# Transcriptional networks are dynamically regulated during cell cycle progression in human Pluripotent Stem Cells

**DOI:** 10.1101/2020.06.14.150748

**Authors:** Anna Osnato, Stephanie Brown, Ludovic Vallier

**Affiliations:** Wellcome – MRC Cambridge Stem Cell Institute, Jeffrey Cheah Biomedical Centre, University of Cambridge, CB2 0AW, Cambridge, UK; Department of Surgery, University of Cambridge, Cambridge, UK

**Author notes:** Corresponding author; current address: BIH Centre for Regenerative Therapies, Charité - Universitätsmedizin Berlin.

## Abstract

Cell cycle progression follows a precise sequence of events marked by different phases and checkpoints which are associated with specific chromatin organisation. Whilst these changes have been extensively studied, their consequences on transcriptional networks remain to be fully uncovered, especially in dynamic model systems such as stem cells. Here, we take advantage of the FUCCI reporter system to show that chromatin accessibility, gene expression and key transcription factors binding change during cell cycle progression in human Embryonic Stem Cells (hESCs). These analyses reveal that core pluripotency factors such as OCT4, NANOG and SOX2 but also chromatin remodelers such as CTCF and RING1B bind the genome at specific phases of the cell cycle. Importantly, this binding pattern allows differentiation in the G1 phase while preserving pluripotency in the S/G2/M. Our results highlight the importance of studying transcriptional and epigenetic regulations in the dynamic context of the cell cycle.

## Introduction

Cell cycle regulation is essential to control the number of cells generated during development and organ homeostasis, while dysregulation of proliferation can result in disease. Importantly, recent studies have suggested that cell cycle regulation could go beyond its canonical function in controlling cell division initiation. Indeed, chromatin alternates between an invariant mitotic conformation and a cell-type specific interphase organisation during cell cycle progression^1^. Loops and Topological-Associated Domains (TADs) insulation emerge very early in the G1 phase, but A/B compartments reappear only during the S phase^2^. Global chromatin decondenses in the G1 phase, and refolds during S/G2/M^3^. In addition, cell fate propensity has been shown to be determined by the cell cycle state of hESCs via the regulation of SMAD2/3, the main effector of the TGFβ signalling pathway^4^. However, the genome wide impact of these changes on transcriptional networks and the link with chromatin structure are still unknown. To address these questions, we used a FUCCI-hESC reporter line^4,5^ to perform ATAC-seq, RNA-seq and ChIP-seq on specific transcription factors (TFs) during cell cycle progression to look at the dynamic changes in chromatin accessibility and transcription factor binding. We observed that the genomic location of several pluripotency factors and epigenetic modifiers are cell cycle regulated, and this allows induction of differentiation in the G1 phase.

## Results

### Chromatin accessibility changes during cell cycle progression in hESCs

FUCCI-hESCs^4^ were grown in chemically defined conditions maintaining pluripotency and then sorted based on their fluorescence (Early G1 (EG1), Late G1 (LG1), G1/S and S/G2/M) (Fig. 1a, Supplementary Information Fig. 1a, b). ATAC-seq (Fig. 1b, Supplementary Information Fig. 1c) was performed and a conservative set of peaks was generated using the IDR pipeline^6^. 88,479 reproducible peaks were identified in EG1, 117,283 in LG1, 99,109 in G1/S, 80,398 in S/G2/M (Supplementary Information Fig. 1d). This suggests that chromatin shows a high degree of accessibility in all cell cycle phases in hESCs especially when compared to definitive Endoderm cells (dEN^7^). Interestingly, the number of open regions was higher in the early cell cycle while slightly decreasing during the S/G2/M phase (Fig. 1c), thereby confirming that chromatin could narrowly close during progression towards cell division, as shown in two key regions upstream the pluripotency genes *POU5F1* and *NANOG* (Fig. 1d). In total, we could identify 153,007 unique open regions. However, only ~33% of these regions were opened in all the phases (50,461) whereas ~67% of these regions were accessible in either a single phase (~34%) or more than one, but not all (~33%) (Fig. 1e). Conservative pairwise comparisons^8^ detected about 1,000 highly dynamic sites (|FC| >1.5 and corrected p-value p-adj < 1E-6), mostly changing through the EG1 phase: 715 regions from EG1 to LG1, 195 from S/G2/M to EG1 (Fig. 1f, g). These observations suggest that chromatin accessibility dynamically changes during cell cycle progression. Interestingly, regions opening in the EG1 phase were enriched for genes associated with transcriptional regulation, whereas regions closing from S/G2/M to EG1 were enriched for genes related to cell cycle (Fig. 1h, Supplementary Information Fig. 1e). This suggests that rewiring of the transcriptional networks required to establish cell identity could occur in the early phase of cell cycle, whereas mechanisms controlling cell division are induced later. Interestingly, changes around promoter regions are most frequent in the EG1 phase, confirming that chromatin around promoters rapidly reorganises after division to regulate transcription (Fig. 1i) thereby suggesting that the EG1 phase could be the most dynamic in terms of transcriptional regulation. In order to understand which transcription factors are regulating such changes, we performed digital footprinting analyses on the regions showing dynamic chromatin accessibility during cell cycle progression. Combining FLR^9^ and Wellington^10^ pipelines, we generated a conservative list of 1,807 footprints. Of these, 74% (1,330) were found in EG1, 8% (147) in LG1, 11% (203) in G1/S, and 7% (127) in S/G2/M (Supplementary Information Fig. 1f). In all the phases, apart from the G1/S transition, the top-ranking footprint corresponded to the transcription factor CTCF (Fig. 1j, k; Supplementary Information Fig. 1g, Supplementary Table 1). More specifically, CTCF represented ~50% of all the footprints in the EG1 phase, dropping to ~10% in the other phases. A wider distribution of factors’ footprints could be detected in the rest of the cell cycle, with a gradual increase in FOX-protein footprints (FOXO3, FOXO4) which were previously described as pioneering factors able to bind condensed chromatin^11^. These data suggest that transcriptional networks are dynamically regulated during cell cycle progression in hESCs and that CTCF could play a role in chromatin regulations especially at the start of the cell cycle.

**Figure 1.**
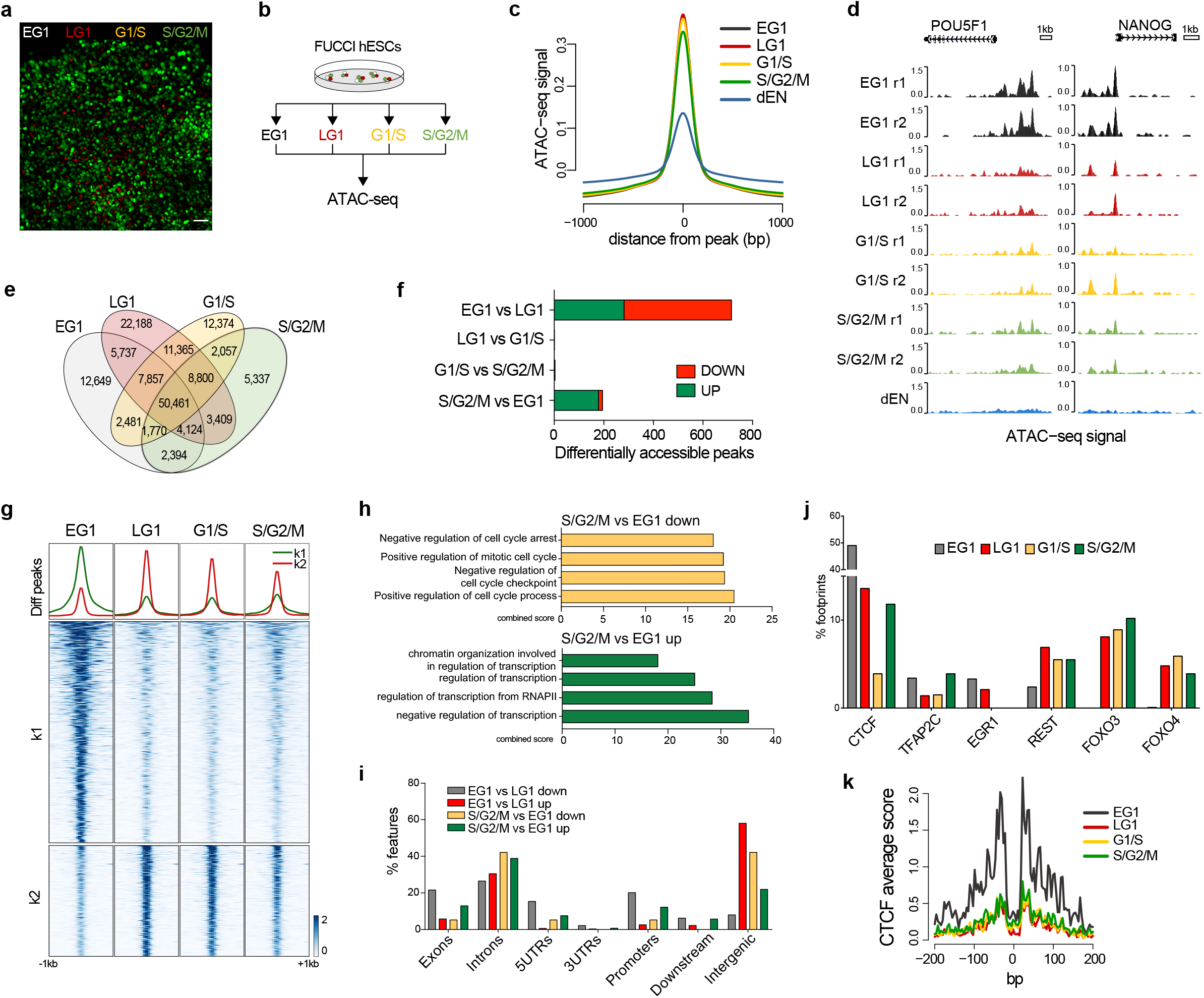
Chromatin accessibility changes during cell cycle progression in hESCs. (**a**) FUCCI hESCs colony showing reporters for the 4 different cell cycle phases: Early G1 (EG1, no fluorescence), Late G1 (LG1, red), G1/S transition (G1/S, yellow), and S/G2/M (green). Scale bar 50μm. (**b**) Experimental approach: FUCCI hESCs were sorted in the 4 cell cycle phases and ATAC-seq was performed on biological duplicates. **(c)** Normalised ATAC-seq signal ±1kb around nucleosome free regions in EG1, LG1, G1/S, S/G2/M, compared to definitive Endoderm (dEN). **(d)** Genome browser tracks reporting chromatin accessibility (ATAC-seq) in cell cycle sorted hESCs at the *POU5F1* and *NANOG* loci in biological duplicates. **(e)** Venn diagram showing ATAC-seq peaks overlap in all 4 cell cycle phases. **(f)** Number of differentially accessible peaks during cell cycle progression obtained by pairwise comparison (|FC| > 1.5, corrected p-value (p-adj) < 1E-6) and their distribution up vs down. **(g)** Heat-maps displaying normalised accessibility signals of differentially accessible regions from (f) averaged using 50bp bins. Regions are centred on the merged peaks for the two replicates ±1kb. K-means clustering corresponds to k=2. **(h)** Gene ontology for Biological Process of differentially accessible peaks in S/G2/M vs EG1 down, S/G2/M vs EG1 up (diffReps, Fig. 1f). Combined score is calculated by multiplying the unadjusted p-values with the z-score. **(i)** Feature distribution (expressed as percentage) for the ATAC-seq peaks for differentially open regions (diffReps) in EG1vsLG1 down, EG1vsLG1 up, S/G2/M vs EG1 down, S/G2/M vs EG1 up. **(j)** Footprints frequency (expressed as percentage) related to single transcription factors in EG1, LG1, G1/S, S/G2/M. **(k)** Average profile of CTCF footprint in the different cell cycle phases generated from Tn5 insertion frequencies.

### Changes in chromatin accessibility during cell cycle are not associated with transcriptional variations

We next decided to assess whether changes in chromatin accessibility correlate with variations in gene expression. Therefore, RNA-seq was performed on FUCCI-hESCs sorted in the different cell cycle phases (EG1, LG1, S/G2/M; Fig. 2a and Supplementary Information Fig. 2a, b). Key pluripotency markers such as *POU5F1* and *NANOG* did not show cell phase specific expression, whereas several cell cycle regulators such as CDK1 showed the expected pattern on expression (Fig. 2b; Supplementary Information Fig. 2c). Interestingly, in agreement with previous reports^12^, some developmental regulators such as *SOX17* and *GATA6* showed higher background expression in the LG1 phase. Differential expression analysis (DESeq2^13^) revealed 394 genes showing cell cycle regulated expression (Fig. 2c, d). Gene Ontology and pathway analyses showed enrichment for cell cycle terms especially regulators of cell division (Fig. 2e). Hierarchical clustering identified three distinct patterns of expression (Fig. 2b, f). Cluster 1 included genes mostly induced during the S/G2/M phase such as *CDK1, CDC25C, AURKA* (Supplementary Information Fig. 2c). Cluster 3 included a list of metabolic genes downregulated from EG1 to S/G2/M, suggesting that metabolic activity might decrease during cell cycle progression. In agreement with our finding of highly dynamic chromatin around the EG1 phase, this cluster included several histone genes clusters, and pathway analysis showed enrichment for chromatin rearrangement terms such as promoter opening and histones acetylation/deacetylation (Supplementary Information Fig. 2d). Cluster 2 includes genes upregulated in LG1, some of which are related to endoderm formation and developmental processes, including *EOMES, MIXL1, SOX17, FOXA2, GATA4* and *GATA6.* In parallel, to make sure that steady-state transcriptomic analysis was representative of subtle changes in dynamic expression, we performed RT-qPCR for key genes using intronic primers in order to interrogate nascent transcription events. This confirmed a similar pattern for developmental regulators as identified by RNA-seq (Fig. 2g; Supplementary Information Fig. 2e, f). However, their expression level remains low when compared to induction observed during endoderm differentiation, suggesting priming of expression rather than full induction. In order to assess whether this transcriptional background correlates with changes in chromatin accessibility, we overlapped all of the 13,478 expressed transcripts (log_2_RPM > 1) with all the associated features of the ATAC-seq peaks: ~85% of expressed genes overlap with accessible regions (Fig. 2h), confirming that most transcribed genes are indeed located in regions of open chromatin. We then overlapped the list of differentially expressed genes with the list of dynamic ATAC regions (from Fig. 1f). Strikingly, only ~3% of differentially expressed genes overlapped with dynamic regions thereby indicating that the majority of cell cycle regulated transcripts originate from constantly open chromatin regions (Fig. 2i; Supplementary Information Fig. 2g). Taken together, these results suggest that developmental regulators are expressed at low level in the G1 phase, potentially marking the priming of hESCs for differentiation. However, this transcriptional “leakiness” originates mostly in constantly open chromatin regions suggesting that it is not induced by changes in accessibility but rather other mechanisms such as differential binding of transcription factors.

**Figure 2.**
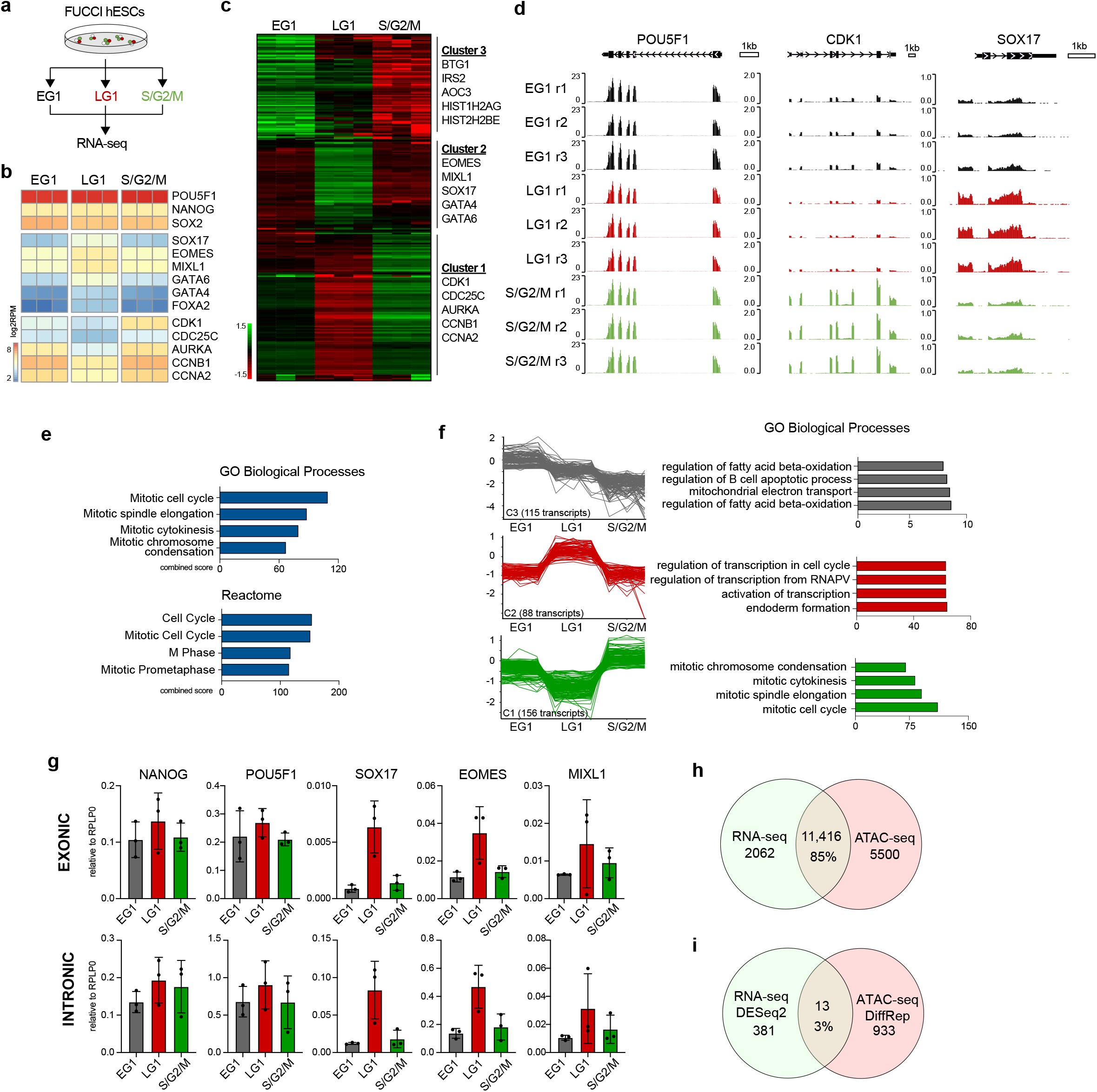
Changes in chromatin accessibility during cell cycle are not associated with transcriptional variations. **(a)** Experimental approach. FUCCI hESCs were sorted in 3 different cell cycle phases: Early G1 (EG1), Late G1 (LG1), G1/S transition (G1/S), and S/G2/M. RNA-seq was performed following sorting on biological triplicates. **(b)** Heatmap reporting log2RPM expression of selected genes (pluripotency markers, developmental regulators, cell cycle) in three biological replicates. **(c)** Hierarchical clustering of 394 differentially expressed genes during cell cycle progression (p-value < 0.05, per probe normalised). Three main clusters and relative representative genes are reported. **(d)** Genome browser tracks reporting cell cycle regulated expression (RNA-seq) in sorted hESCs of *POU5F1, CDK1* and *SOX17* in 3 biological replicates. **(e)** Gene Ontology (Biological Processes and Reactome) for the 394 differentially expressed genes ranked by combined score. Combined score is calculated by multiplying unadjusted p-value with z-score. **(f)** Average expression and related gene Ontology (Biological Processes) for the three main clusters in (c), ranked by combined score. Combined score is calculated by multiplying unadjusted p-value with z-score. **(g)** RT-qPCR expression analysis of cell cycle sorted FUCCI hESCs using standard primers spanning exon-exon junctions (exonic) and primers designed over introns (intronic) to measure pre-processed transcripts for pluripotency and genes selected from (c). Expression is relative to the housekeeping gene *RPLP0*. **(h)** Venn diagram summarising overlap of expressed genes (RNA-seq, log_2_RPM > 1) and accessible chromatin (ATAC-seq). 85% represents the percentage of expressed genes present in open chromatin regions (overlapping ATAC peaks). **(i)** Venn diagram reporting overlap of differentially expressed genes (RNA-seq DESeq2) and regions changing accessibility (ATAC-seq DiffRep). 3% represents the percentage of differentially expressed genes overlapping with changes in chromatin accessibility.

### Transcription factors binding changes during cell cycle progression in hESCs

In order to test whether transcription factors are driving these transcriptional changes, we performed ChIP-seq in sorted FUCCI-H9 for different TFs (Fig. 3a). We first looked at CTCF binding in order to validate footprinting predictions. Genome wide mapping revealed 5,606 unique binding sites, of which 4,401 were found in EG1 confirming its higher occupancy in the early cell cycle (Fig. 3b-d). This pattern was confirmed looking at previously identified key loci^14^ such as the *LEFTY/MIXL1,* where CTCF anchor sites that regulate this super enhancer are strongly present in the EG1 phase, and gradually decrease during cell cycle progression (Fig. 2e). Interestingly, this locus also includes the *MIXL1* and *H3F3A* genes (Fig. 2f), the latter being the top hit for CTCF differential binding analysis (Supplementary Information Fig. 3a, Supplementary Table 2). *H3F3A* encodes for the histone H3.3, previously described as being enriched at the centre of the intergenic CTCF-binding sites^15^, necessary for correct cell cycle progression and to regulate heterochromatin structures^16^. Of the 708 CTCF footprints identified using ATAC-seq data ~60% overlapped with CTCF ChIP-seq confirming the robustness of our prediction (Supplementary Information Fig. 3b). Interestingly, 77% of CTCF binding sites were located in open chromatin regions, with a higher overlap in the EG1 phase (Supplementary Information Fig. 3c), thereby suggesting a specific role for CTCF in regulating chromatin accessibility in this phase of the cell cycle. All together, these results show that CTCF genome wide occupancy changes during cell cycle progression. The increased binding of CTCF in EG1 supports the hypothesis that CTCF could drive chromatin restructuring to regulate transcription^17^ and genome integrity^18^ during the subsequent cell cycle.

**Figure 3.**
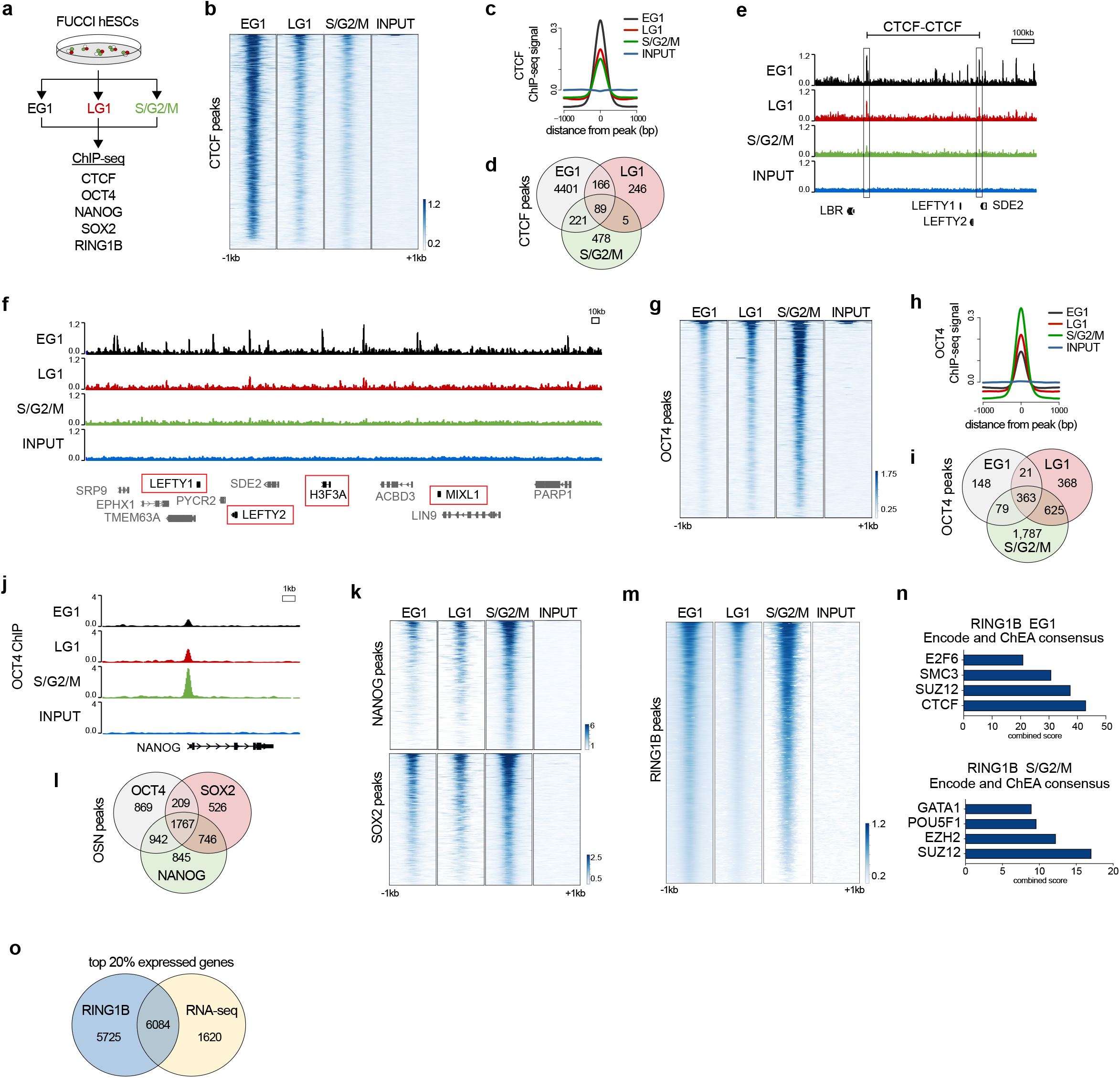
Transcription factors binding changes during cell cycle progression in hESCs. (**a**) Experimental approach. FUCCI hESCs were sorted in 3 different cell cycle phases: Early G1 (EG1), Late G1 (LG1), and S/G2/M. ChIP-seq was performed following sorting on biological duplicates for CTCF, OCT4, NANOG, and SOX2. **(b)** Heatmaps displaying CTCF binding in the different cell cycle phases EG1, LG1, S/G2/M, compared to INPUT. **(c)** Normalised ChIP-seq signal ±1kb around all peaks in EG1, LG1, S/G2/M, compared to INPUT. **(d)** Venn diagram reporting overlap of CTCF binding sites during cell cycle. **(e)** Genome browser tracks reporting CTCF binding in cell cycle sorted hESCs at the *LEFTY* locus. A selected CTCF-CTCF loop^14^ is depicted as a black line. A subset of genes present in this loop is shown for simplicity. **(f)** Genome browser tracks reporting CTCF binding in cell cycle sorted hESCs at the *LEFTY/H3F3A/MIXL1* adjacent loci. Relevant genes are highlighted in red. **(g)** Heatmaps displaying OCT4 binding in the different cell cycle phases EG1, LG1, S/G2/M, compared to INPUT. **(h)** Normalised OCT4 ChIP-seq signal ±1kb around all peaks in EG1, LG1, S/G2/M, compared to INPUT. **(i)** Venn diagram showing peaks overlap for OCT4 ChIP-seq in EG1, LG1, S/G2/M. **(j)** Genome browser tracks reporting OCT4 binding in cell cycle sorted hESCs at the *NANOG* promoter. **(k)** Heatmaps displaying NANOG and SOX2 binding in the different cell cycle phases EG1, LG1, S/G2/M, compared to INPUT. **(l)** Venn diagram showing binding sites overlap for OCT4, NANOG and SOX2. **(m)** Heatmaps displaying RING1B binding in the different cell cycle phases EG1, LG1, S/G2/M, compared to INPUT. **(n)** Gene ontology for Encode ChEA consensus sequences of RING1B EG1 and S/G2/M consensus sequences. Combined score is calculated by multiplying the unadjusted p-values with the z-score. **(o)** Overlap of RING1B bound genes with highly expressed genes by RNA-seq (genes are ranked based on their expression, and the top 20% are selected). For all the heat-maps, regions are centred on the merged peaks for the two replicates ±1kb in all the phases and compared to INPUT.

In order to check whether this binding pattern could be shared with other TFs or it is uniquely related to the role of CTCF in the early cell cycle, we performed ChIP-seq on sorted FUCCI-H9 for the core pluripotency factors OCT4, NANOG and SOX2 (OSN). Interestingly, we found a different binding pattern shared between the three factors with higher genome occupancy in the late cell cycle. For OCT4, we found a total of 4,843 binding sites distributed as follows: 611 in EG1, 1,378 in LG1, and 2,854 in S/G2/M (Fig. 3g-i) showing that the number of genomic regions bound by this key pluripotency factor increases during cell cycle progression, as shown for the *NANOG* locus (Fig. 3j). The same binding pattern was observed for NANOG and SOX2 with both factors binding the genome more frequently in the S/G2/M phase (Fig. 3k; Supplementary Information Fig. 3d, e). OSN combinatorial analysis (Fig. 3l) showed that of the 5,904 unique peaks, 30% (1,767) were shared, with the majority involved in signalling pathways regulating pluripotency (Supplementary Information Fig. 3g). Single factors revealed specific features: OCT4 peaks were enriched for deubiquitination terms, NANOG for insulin receptor signalling pathway, and SOX2 involved in neuron differentiation.

Intrigued by these findings, we asked whether additional factors known to be linked to the core transcriptional network might behave similarly. For that we performed ChIP-seq for RING1B, a chromatin modifier shown to partially bind chromatin in an OCT4-dependent manner^19^. These analyses revealed a unique binding pattern with high occupancy in the EG1 phase, a decrease in the LG1, and an increase in the S/G2/M (Fig. 3m, Supplementary Information Fig. 3f). Interestingly, RING1B is known to have several roles from controlling chromatin compaction to regulating gene expression^20^. Thus, our observations could imply that these divergent roles might have a temporal distribution during cell cycle progression. Of all the bound sites, gene ontology analysis for Biological Process highlighted its role in regulating transcription (Supplementary Information Fig. 3h). Interestingly, regions bound by RING1B in EG1 were enriched for CTCF binding sites together with SMC3, a core subunit of the cohesin complex (Fig. 3n), suggesting that RING1B might have a structural role in opening chromatin at the start of the cell cycle. However, pluripotency factors such as OCT4 are enriched in S/G2/M (Fig. 3n), validated by the stronger binding of OCT4 at binding sites shared with RING1B in S/G2/M (Supplementary Information Fig. 3i). To further validate this dual role at the transcriptional level, we then overlapped RING1B peaks with top 20% expressed genes, and found that ~50% of the genes bound by RING1B are highly expressed (Fig. 3o), reinforcing the hypothesis that RING1B does not only have a repressive role but it is also bound to highly transcribed genes. Taken together, these findings show that transcription factors have different binding patterns during cell cycle progression and that this regulation could define their activity.

The dynamic binding of these transcription factors could be explained by a variety of mechanisms including variation in their expression during cell cycle progression. However, RNA-seq and RT-qPCR validations showed that the expression of these genes does not change between cell cycle phases (Fig. 4a, b). For this reason, we looked at the protein level of these TFs in sorted FUCCI-H9. Pluripotency factors’ expression levels were quite variable and did not show a clear expression pattern (Fig. 4c, d; Supplementary Information Fig. 4a). Therefore, variations in the protein levels cannot explain their differential binding. Interestingly, CTCF instead showed a trend of higher expression in the early cell cycle, in line with its chromatin binding (Fig. 4e, f), suggesting that different kinds of regulation are present during cell cycle progression. For pluripotency factors instead, we hypothesised that post-translational modifications could regulate their differential binding. Of particular interest is the phosphorylation at Serine 236 of OCT4 (pOCT4) which is known to regulate OCT4 binding to chromatin and its transcriptional activity in hESCs^21^. This modification is known to be the equivalent phosphorylation of mouse Serine 229, which sterically hinders DNA binding^22^. To confirm that such a mechanism could control OCT4 binding in EG1, we performed immunostaining for pOCT4 (Ser236) in FUCCI-hESCs. These analyses showed that mitotic cells have enriched phosphorylation signal, thereby confirming findings in mouse Embryonic Stem Cell (mESCs)^23^. However, hESCs in EG1 still partially retain pOCT4 (Fig. 4g; Supplementary Information Fig. 4b, c; white circles), suggesting that complete dephosphorylation only happens past the EG1 phase. This was further validated with flow cytometry, showing that a similar fraction of cells retains pOCT4 in S/G2/M and EG1 compared to unsorted cells and LG1 (Fig. 4h, i; Supplementary Information Fig. 4d). Taken together, these results suggest that phosphorylation could regulate the genomic binding of OCT4 during cell cycle progression also in hESCs. Importantly, EG1 has been shown to be a key phase during cell cycle for cell fate decision in hESCs^4^. For this reason, we hypothesised that the reduced capacity of pluripotency factors to bind the genome in EG1 could be necessary for induction of differentiation.

**Figure 4.**
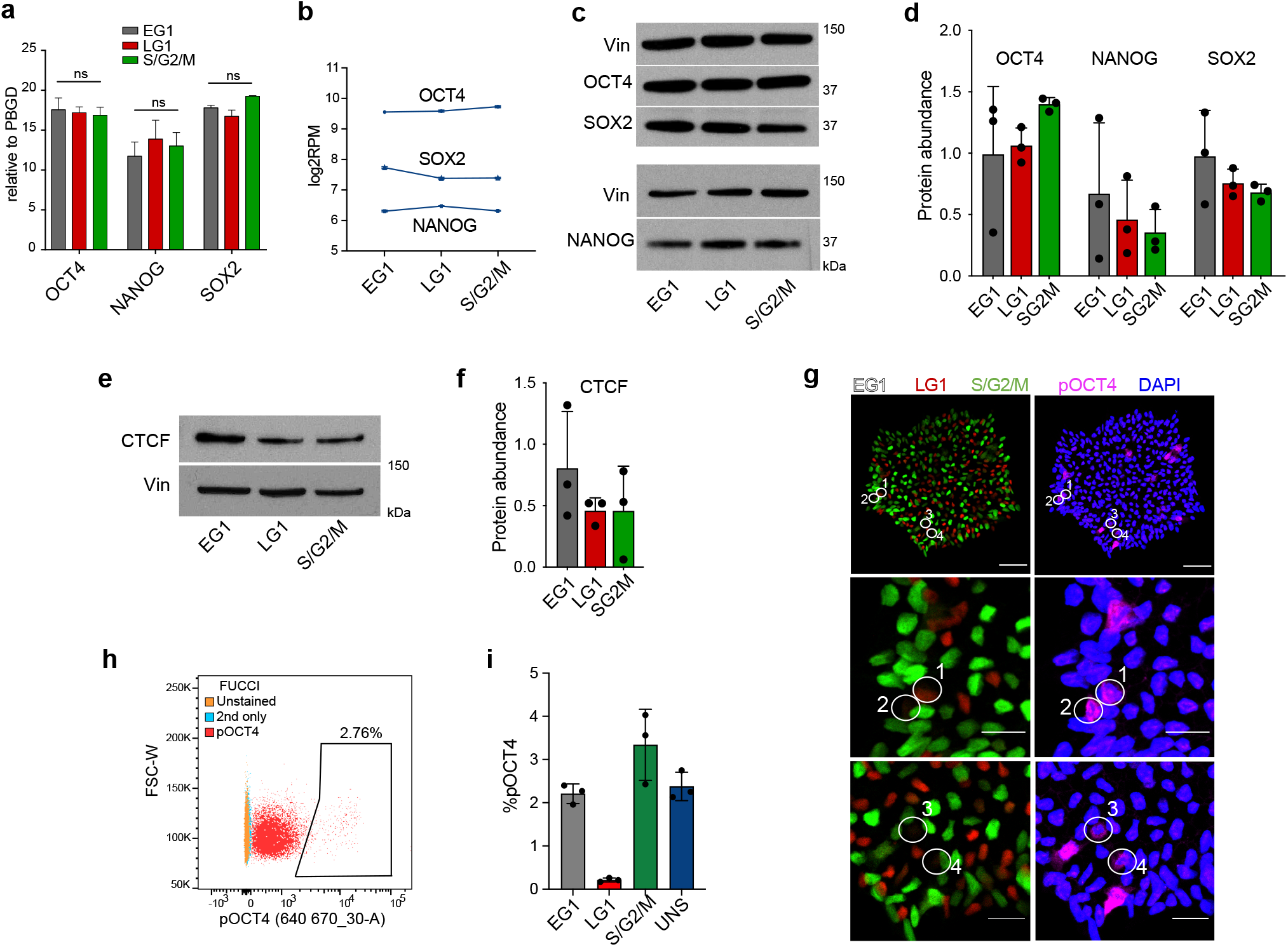
Post-translational modification of OCT4 regulates its chromatin binding. **(a)** RT-qPCR for *OCT4, NANOG* and *SOX2* in EG1, LG1, and S/G2/M relative to housekeeping gene *PBGD* (two-ways ANOVA, n=3). **(b)** RNA-seq data for *OCT4, SOX2*, and *NANOG*, in EG1, LG1, and S/G2/M, expressed in log_2_RPM. **(c)** Representative western-blot for OCT4, SOX and NANOG in sorted EG1, LG1, and S/G2/M, plus vinculin (Vin) as loading control. **(d)** Relative quantification of (c) performed using the software Fiji (ImageJ) reporting OCT4/SOX2/NANOG to vinculin ratio. Each dot represents a biological replicate (n=3). **(e)** Representative western-blot for CTCF in sorted EG1, LG1, and S/G2/M, plus vinculin (Vin) as loading control. **(f)** Relative quantification of (e) performed using the software Fiji (ImageJ) reporting CTCF to vinculin ratio. Each dot represents a biological replicate (n=3). **(g)** Immunofluorescence for phospho-OCT4 Ser236 (pOCT4, magenta) and nuclear staining (DAPI, blue) in FUCCI hESCs (S/G2/M cells in green-mAG, LG1 cells in red-mKO2, EG1 cells no colour). Bottom two panels report higher magnification of numbered circled cells in the EG1 phase. Scale bars: top panel 50μm, bottom panels 20μm. **(h)** Flow cytometry plots of FUCCI hESCs positive for pOCT4 in all cell cycle phases. Unstained cells and secondary (2nd only) reported as negative controls. **(i)** Quantification (expressed as percentage) of sorted FUCCI hESCs positive for pOCT4 divided by cell cycle phase compared to the unsorted control.

### Cell cycle specific OCT4 binding is required for induction of differentiation

To test whether pluripotency factor binding regulates initiation of differentiation, we developed two FUCCI-hESC lines to respectively over-express or knock-down OCT4 in a cell cycle phase specific manner (Fig. 5a). For that, we used a tamoxifen (4OHT) inducible form of OCT4 (OCT4-ERT2), that relies on the fusion of OCT4 with a modified fragment of the Estrogen receptor^24^ (Supplementary Information Fig. 5a). Western blot confirmed correct integration (Supplementary Information Fig. 5b) and immunostaining confirmed correct nuclear translocation of OCT4 upon addition of 4-OHT in as little as 4h in both pluripotent and neuroectoderm cells (Fig. 5b). We then sorted OCT4-ERT2-FUCCI-hESCs in EG1 in order to start the differentiation when OCT4 is not bound to chromatin, and re-plated them in culture condition inducing endoderm differentiation^25,26^ for 24h in the presence or absence of 4-OHT for the first 4h. Interestingly, OCT4 induction decreases the expression of the key early endoderm markers such as *T*, *MIXL1* and *GSC* (Fig. 5c) suggesting a less efficient differentiation. In parallel, we generated an inducible hESCs for the knockdown of OCT4 expression (iKD-OCT4-FUCCI-hESCs), taking advantage of the Tetracycline (TET) based system OptiKD^27^ (Fig. 5d; Supplementary Information Fig. 5c, d). iKD-OCT4-FUCCI-hESCs sorted in S/G2/M were re-plated in culture condition inducing endoderm differentiation for 24h in presence or absence of TET, in order to deplete OCT4 when its binding is at its peak during the cell cycle. Interestingly, decrease of OCT4 in the S/G2/M phase promoted the expression of late endoderm (*SOX17*) and mesoderm (*HAND1*) markers (Fig. 5e) thereby suggesting a more efficient differentiation process. Taken together, these results show that variation in OCT4 binding during cell cycle is required for optimal cell fate decision in hESCs or more broadly, that cell cycle specific regulation of transcriptional networks could control capacity of differentiation in hESCs.

**Figure 5.**
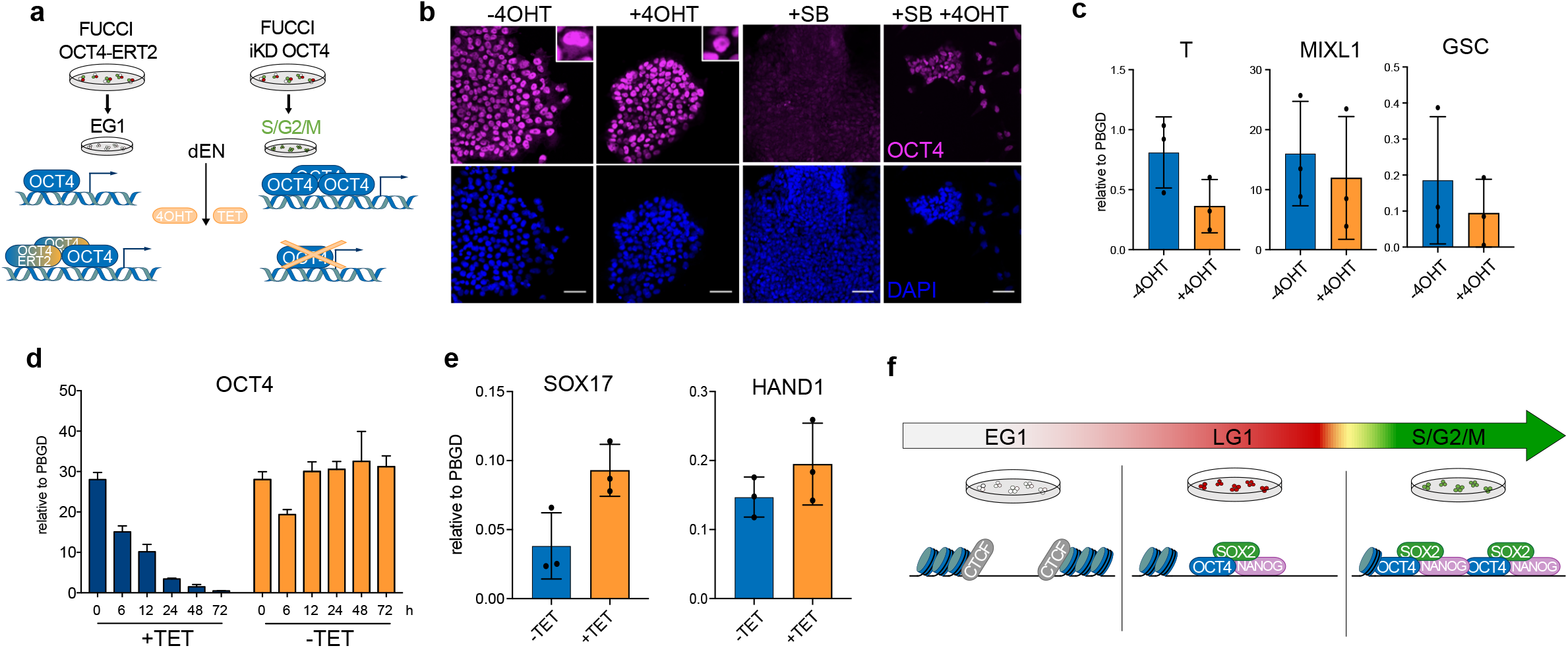
Cell cycle specific OCT4 binding is required for induction of differentiation. **(a)** Experimental approach: FUCCI OCT4-ERT2 were sorted in Early G1 (EG1) when OCT4 binding is at its minimum and re-plated in Endoderm inducing conditions (dEN). OCT4 nuclear translocation was induced upon 4-Hydroxytamoxifen addition (4OHT). FUCCI iKD OCT4 were sorted in S/G2/M when OCT4 binding is at its max and re-plated in dEN conditions. OCT4 knock-down was induced upon tetracycline addition (TET). **(b)** Immunofluorescence for OCT4 (magenta) and nuclear staining DAPI (blue) in FUCCI OCT4-ERT2. In presence of 4-Hydroxytamoxifen (+4OHT), OCT4 is uniquely nuclear. Upon neuroectoderm induction (SB), endogenous OCT4 is strongly reduced and upon 4-Hydroxytamoxifen induction is translocated into the nucleus (SB +4OHT). Scale bar 50μm. **(c)** RT-qPCR expression of endoderm markers *T*, *MIXL1*, and *GSC* in presence or absence of 4 hours of 4-Hydroxytamoxifen (+4OHT) in EG1 sorted cells after 24h of differentiation. Expression reported as relative to housekeeping gene *PBGD* **(d)** RT-qPCR reporting OCT4 expression time-course (0-72 hours) during knock-down in presence of tetracycline (+TET) compared to -TET control. Expression relative to housekeeping gene *PBGD*. **(e)** RT-qPCR reporting expression of late endoderm markers *SOX17* and *HAND1* in presence or absence of tetracycline (TET) in S/G2/M sorted cells after 24h of differentiation. **(f)** Schematic summary of the proposed working model. In the EG1 phase, immediately after cell division, chromatin is highly dynamic and CTCF binding is proposed to regulate changes in its accessibility. Core pluripotency factors OCT4, SOX2 and NANOG (OSN) show low occupancy in the G1 phase, and this allows hESCs to respond to differentiation cues. In the S/G2/M phase, core pluripotency factors fully re-bind chromatin, re-establishing the pluripotency network while sustaining self-renewal and blocking differentiation.

## Discussion

Our study investigates for the first time the dynamic mechanisms that coordinate chromatin accessibility, gene expression, and transcriptional networks during cell cycle progression in hESCs. In the EG1 phase, immediately after cell division, chromatin is highly dynamic and CTCF appears to regulate changes in DNA accessibility. Indeed, CTCF might be the driver for reorganising chromatin after chromosome condensation during cell division^17,28^ in conjunction with other chromatin modifiers such as RING1B^29^. In parallel, the core pluripotency factors OCT4, SOX2 and NANOG show low occupancy in the G1 phase allowing hESCs to rapidly respond to differentiation cues. In the S/G2/M phase, they fully bind chromatin to re-establish the pluripotency network, sustain self-renewal and block differentiation. These results reinforce previous studies showing that this phase actively promotes the pluripotent state^30^. This change in binding might be regulated via post-translational modifications such as phosphorylation, preventing OCT4 from binding DNA in EG1^21^. Importantly, these changes could define the temporal function of RING1B which might help CTCF in restructuring chromatin in EG1 while acting as transcriptional regulator in S/G2/M to maintain pluripotency^31^. Ultimately, these cell cycle specific interplays are necessary to coordinate induction of differentiation and exit of pluripotency with cell division.

Our results also demonstrate that key transcription factors are cell cycle regulated and dynamically change their genomic location. This binding pattern suggests that transcription factor activity and specificity are coordinated with cell cycle progression. This process could represent a key mechanism that allows stem cells to not only maintain their capacity to change cellular identity but also for transcription factors to have different functions based on the cell cycle state of the cells. Importantly, these findings highlight the importance of studying transcriptional and epigenetic mechanisms in the context of the cell cycle. More generally, most ChIP-seq studies are currently capturing transcription factors’ binding on the most extended cell cycle phase which can vary from cell to cell. Thus, we would like to propose that genomic location of transcription factors should be investigated in a cell cycle dependent manner, since phase-specific analyses could reveal new molecular functions even for commonly studied regulators. This novel concept could apply to any cell type and thus have a major impact on our understanding of transcription factors’ functions.

## Acknowledgements

The work was supported by the Wellcome PhD programme in Stem Cell Biology (A.O.), the European Research Council starting grant “Relieve IMDs” (L.V.); and a core support grant from the Wellcome and Medical Research Council to the Cambridge Stem Cell Institute. We thank Dr Siim Pauklin for providing the ATAC-seq samples. We thank the Cambridge NIHR BRC Cell Phenotyping Hub for help with cell sorting and the DNA Pipelines Operations at the Wellcome Sanger Institute for next generation sequencing. We also thank Professor Hong-Duk Youn and Jihoon Shin for kindly providing pOCT4 antibody.

## Author contributions

A.O. conceived the study, performed all the experiments, analysed data and wrote the manuscript. S.B. helped with experimental planning, data analysis and reviewed the manuscript. L.V. conceived, supervised, and supported the study, wrote and provided final approval of the manuscript.

## Methods

### hESCs culture and germ layers specification

FUCCI-hESCs (H9/WA09 line; WiCell) were grown feeder- and serum-free as previously described^32,33^. Briefly, cells were grown in either in-house E6 supplemented with 25ng/ml Fibroblast growth factor 2 (FGF2) and 2ng/ml TGFβ (BioTechne) or in Chemically Defined Medium (CDM), supplemented with Bovine Serum Albumin (BSA), recombinant Activin-A (10 ng/ml) and 12 ng/ml FGF2 on 10μg/ml Vitronectin-XF (STEMCELL Technologies) or 0.1% Gelatin and MEF-medium coated plates, respectively (Supplementary Table 3). If not specified, growth factors were provided by Dr Marko Hyvonen (Department of Biochemistry, University of Cambridge). Cells were passaged weekly using either EDTA or Collagenase IV and plated as clumps of 50-100 cells dispensed at a density of 100-150 clumps/cm^2^. All the FUCCI-hESCs lines used were kept under selection with Puromycin (1mM) and Geneticin (G-418, 0.2 mg/ml), or alternatively G-418 (0.2 mg/ml) and Blasticidin (1mM). Definitive Endoderm (dEN) was induced for 3 days in CDM-Polyvinyl alcohol (PVA) without insulin supplemented with 20ng/ml FGF2, 10μM LY294002, 100ng/ml Activin A, and 10ng/ml BMP4, changing media daily, as previously described^25,26^. Neuroectoderm was induced for 6 days in CDM-BSA supplemented with 12ng/ml FGF2 and 10μM SB431542 (Activin/Nodal/TGFβ inhibitor; Tocris), as previously described^34^. hESCs were monthly monitored for mycoplasma infection.

### Immunofluorescence (IF)

Cells were grown on glass coverslips coated with either Vitronectin XF or Gelatine and fixed with 4% PFA for 10 min at RT, rinsed twice with PBS, and permeabilised for 20 min at RT using PBS/0.25% Triton X-100 (Sigma). Cells were blocked for 30 min at RT with blocking solution (PBS-0.25% Triton X-100 plus BSA 1%). Primary and secondary antibodies (listed in Supplementary Table 4) were diluted in blocking solution and incubated for 1 h at 37 °C. Cells were washed twice with blocking solution after each antibody staining, and stained with DAPI for 5 min at RT (0.1 mg/ml DAPI in PBS-0.1% Triton). Finally, coverslips were mounted on slides using ProLong Glass antifade reagent (ThermoFisher Scientific) and imaged using an LSM 700 confocal microscope (Zeiss). Images were processed using the software Fiji (ImageJ). At least four different fields from each experiment were imaged and representative ones are shown in the figures.

### Western Blotting

For whole cell lysates, cells were washed once in D-PBS and resuspended in ice cold RIPA buffer (150mM NaCl, 50mM Tris, pH8.0, 1% NP-40, 0.5% sodium deoxycholate, 0.1% sodium dodecyl sulfate) containing protease and phosphatase inhibitors for 10 min. Protein concentration was quantified by a BCA assay (Pierce) following the manufacturer’s instructions using a standard curve generated from BSA and read at 600 nm on an EnVision 2104 plate reader. Samples were prepared by adding 4x NuPAGE LDS sample buffer (ThermoFisher Scientific) plus 1% b-mercaptoethanol and heated at 95 °C for 5 min. 5–10mg of protein per sample was run on a 4–12% NuPAGE Bis-Tris Gel (ThermoFisher Scientific) and then transferred to PVDF membrane by liquid transfer using NuPAGE Transfer buffer (ThermoFisher Scientific). Membranes were blocked for 1 hr at RT in PBS 0.05% Tween-20 (PBST) supplemented with 4% non-fat dried milk and incubated overnight at 4 °C with primary antibodies diluted in the same blocking buffer (Supplementary Table 4). After three washes in PBST, membranes were incubated for 1 hr at RT with horseradish peroxidase (HRP)-conjugated secondary antibodies diluted in blocking buffer, then washed a further three times before being incubated with Pierce ECL2 Western Blotting Substrate (ThermoFisher Scientific) and exposed to X-Ray Film. Relative quantification was performed using Fiji (ImageJ). All western blots were performed at least in three biological replicates.

### RT-qPCR

RNA was extracted using the GenElute Mammalian Total RNA Miniprep Kit and the On-Column DNase I Digestion Set (both from Sigma-Aldrich) following manufacturer’s instructions. cDNA was synthesised using 500ng of RNA in a reaction with 250ng random primers, 0.5mM deoxynucleotides (dNTPs), 20U RNaseOUT, and 25U of SuperScript II (all Invitrogen) in a total of 20μl. cDNA was diluted 30-fold and 5μl were used for qPCR using SensiMix SYBR low-ROX (Bioline) with 150nM Forward and Reverse primers (Sigma-Aldrich; Supplementary Table 5 for primer sequences). Samples were run in technical duplicates on 96-well plates on a Stratagene Mx-3005P (Agilent) or 384-well plate on a QuantStudio Flex Real-Time PCR System, and results were analysed using the delta-delta cycle threshold (ΔΔCt) approach using PBGD as housekeeping gene. Primers were designed using PrimerBlast (http://www.ncbi.nlm.nih.gov/tools/primer-blast/), and tested to produce a single PCR product, with an efficiency between 80% and 120%.

### Cell Sorting

FUCCI-hESCs were sorted as previously described^4^. Briefly, cells were washed with PBS and collected by incubating with CDB for 10 min at 37°C. When needed, crosslinking of proteins to DNA was performed prior to sorting, according to the specific experiment (see Chromatin Immunoprecipitation). Cells were washed with 1% PBS/BSA and incubated in 1% PBS/BSA with anti-TRA-1-60-antibody mouse monoclonal IgM (1:100, Santa Cruz Biotechnology, Inc.) for 20 min on ice, together with the secondary antibody, Alexa Fluor® 647 goat anti-mouse IgM (1:1,000, Thermo Fisher). Cells were washed with PBS, re-suspended in 6ml of PBS and sorted into the different cell cycle phases using BD FACS AriaTM III, BD Fusion or BD Influx sorters by the Cambridge NIHR BRC Cell Phenotyping Hub. Gating strategy set as following: cells were then size selected for Forward (FS) and Side Scatter (SS), and doublets were removed. TRA-1-60+ cells were then sorted in the four different cell cycle phases according to the FUCCI reporter: Early G1 (EG1) for double negative (mKO2- / mAG-), Late G1 (LG1) for red-mKO2+, G1/S transition for double positive (yellow-mKO2+ / mAG+), and S/G2/M for green-mAG+. Sorted cells were then used for ATAC-seq, ChIP-seq, and RNA-seq. All the experiments have been performed normalising for cell number, usually to the lowest sample, depending on the downstream application.

### Chromatin Immunoprecipitation (ChIP)

Crosslinking of protein to DNA was performed prior to sorting in 1% PBS/Formaldehyde, rocking for 10 min at room temperature. All the experiments were performed with single crosslinking and neutralised with Glycine (0.125M), for 5 min shaking at room temperature. ChIP was then carried out using LowCell# ChIP kit (C0101007, Diagenode) and DNA recovered using iPure kit (C03010012, Diagenode) according to manufacturer’s instructions, with the following modifications. 5μg of antibodies were used per IP, and the quantity of magnetic beads adjusted accordingly, based on their binding capacity (10μl of beads for ~3μg). Elution was made in Buffer C from the iPure kit, compatible with qPCR and next generation sequencing. For all the experiments, the same number of cells was used per IP after sorting, normalised for the less abundant cell cycle phase, usually EG1 ~1M cells.

### Assay for transposase-accessible chromatin (ATAC)

TRA-1-60+ hESCs were sorted in the different cell cycle phases (EG1, LG1, G1/S, S/G2/M). Samples were generated from two independent experiments, 100,000 cells per sample, as previously described^35^ with some modifications. In brief, cells were washed with ice-cold PBS and incubated with 500μl of ice-cold sucrose buffer (10mM Tris-Cl pH 7.5, 3mM CaCl2, 2mM MgCl2 and 0.32M sucrose) for 12 min on ice. To collect the nuclei, Triton X-100 was added to a final concentration of 0.5% and cells incubated for additional 6 min on ice after mixing. Samples were then resuspended in 1ml of sucrose buffer and transferred to a fresh 1.5 ml Eppendorf tube. The nuclei were centrifuged at 1500g for 3 min at 4°C and the supernatant discarded. Nuclei pellets resuspended in 50μl of Nextera DNA Sample Preparation Kit (Illumina FC-121-1030) for tagmentation, comprising 25μl 2x Tagment DNA buffer, 20μl nuclease-free water and 5μl Tagment DNA Enzyme 1. The tagmentation reaction mixture was transferred in a 1.5ml low-binding tube and incubated at 37°C for 30 min and stopped by the addition of 250μl Buffer PB (Qiagen). The tagmented chromatin was purified using the MinElute PCR purification kit (Qiagen 28004), according to the manufacturer’s instructions, eluting in 11.5μl of buffer EB (Qiagen). For PCR amplification, 10μl of the tagmented chromatin was mixed with 2.5μl of Nextera PCR primer cocktail and 7.5μl of Nextera PCR master-mix (Illumina FC-121-1030) in low-binding PCR tubes. Primers were from the Nextera Index kit (Illumina FC-121-1011), using 2.5μl of an i5 primer and 2.5μl of an i7 primer per PCR, totalling 25μl. PCR amplification was performed as follows: 72°C for 3 min and 98°C for 30 sec, followed by 12 cycles at 98°C for 10 sec, 63°C for 30 sec and 72°C for 3 min. The sample volumes were then increased to 50μl by adding Qiagen EB buffer. The PCR primers were removed with 1x 0.9:1 SPRI beads (Beckman Coulter, Cat no. A63880) according to manufacturer’s instructions and samples eluted in 20μl. Following amplification, excess of primers was removed running the samples of 50ml 1% agarose TAE gel at 90V for 25 min, and size selection obtained by cutting from 120bp to 1kb. MinElute Gel Extraction kit (Qiagen 28604) was used to extract DNA, eluted in 20μl of buffer EB.

### Transfection

The generation of the inducible lines was performed using lipofectamine transfection. Lipofection was performed as previously described^36^. In brief, FUCCI-hESCs cells were seeded into 6 well plates using medium without Penicillin-Streptomycin (Pen/Strep, Gibco), in low density. After 48-72h (depending on colony size), cells were transfected using Lipofectamine® 2000 Transfection Reagent (Thermo Fisher), according to manufacturer’s instructions. In sum, the medium was aspirated and replaced with 1mL/well of Opti-MEM® Reduced Serum Media (Thermo Fisher). 4μg of (each) plasmid was resuspended in 0.25ml of Opti-MEM medium/well. 10μl of Lipofectamine 2000 were dissolved in 0.25ml of Opti-MEM medium/well, and incubated for 5 min at room temperature. Lipofectamine 2000 and the plasmid were mixed together and incubated for 20 min at room temperature. 0.5ml of the DNA/Lipofectamine mix was added drop by drop to the well and incubated for 18-20h. Opti-MEM was then replaced with regular media without antibiotics. Antibiotic selection was started 4-5 days after the transfection (Puromycin 1 mg/ml, G-418 50μg/ml). After 6 days, clones were picked into a 24 wells plate. Each clone was passaged into 12 well plates, for analysis of transgene expression by qPCR, immunostaining or western blot.

### Bioinformatics analyses and pipelines

The following data processing pipelines have been used for all the bioinformatics analyses according to ENCODE guidelines^6^.

### ChIP-seq

TRA-1-60+ hESCs were sorted in the different cell cycle phases (EG1, LG1, S/G2/M). From two independent experiments, chromatin immunoprecipitation was performed as described above. Library preparation and sequencing were performed at the Wellcome Trust Sanger Institute next-generation sequencing facility. Size selection was applied, and fragments between 100bp and 400bp (average length of 200bp) were used to prepare barcoded sequencing libraries using NEBNext Sample Prep Kit1 (NEB) following manufacturer’s instructions. 10ng of input material (as a pool of the different phases) for each experiment was processed in parallel as control. To test whether pooled inputs were an acceptable control, one experiment was run with one input per cell cycle phase. No differences were reported with > 90% peaks shared with the pooled input (data not shown). Equimolar amounts of each library were pooled, and 10 samples/lane were multiplexed on Illumina HiSeq 2000, 2 X 75bp paired-end reads. On average, 400M reads per lane were generated, mapped to human genome GRCh38 reference assembly and stored as cram files. Cram were converted in bam keeping only reads with mapping score above 10 (q>10) using Samtools view^37^. From two independent replicates, consistency between replicates was measured using the Irreproducibility Discovery Rate (IDR) pipeline (https://sites.google.com/site/anshulkundaje/projects/idr). This approach is based on an independent peak-calling for the two replicates with a relaxed cut-off (-p1e-1, --to-large) using MACS^38^. This allows to sample enough noise in the experiment for efficient statistical comparison of replicates in subsequent steps, meaning that many false positives are present in this peak set and they are not considered as final peaks for the separate replicates. Peaks are then ranked for p-value, and the same number is compared based on reproducibility (top 100,000 peaks). This is used to score rankings between peaks in each replicate to determine an optimal cut-off for significance. The final peak calls (bed and bigBed format) are the set of peak calls that pass IDR at a threshold of 5-10% max. This is the conservative peak set and it is used for further analyses. If the minimum number of recommended reads is not achieved, replicates are pooled. For visualisation purposes, biological replicates were combined. Normalised bedGraph format files were generated for each sample using BEDTools v2.24.0^39^. The reads mapped at both DNA strands from 5’ to 3’ direction were extended to a length of 300bp, and the read-enrichment was normalised by million mapped reads and size of the library. bedGraph files were converted to bigWig using UCSC tool bedGraphToBigWig, and visualised on the Biodalliance genome viewer^40^. Differential binding analysis was performed using the R package Diffbind^41^. For the annotation of genomic intervals, Heatmaps and Metaplot generation, the R packages Genomation and ChIPpeakAnno were used^42,43^. GREAT (Genomic Regions Enrichment of Annotations Tool) was used to associate genes with regulatory regions^44^, combined with UCSC liftOver tool to convert bed coordinates from Homo Sapiens GRCh38/hg38 assembly to GRCh37/hg37 assembly (https://genome.ucsc.edu/cgi-bin/hgLiftOver).

### ATAC-seq

TRA-1-60+ hESCs were sorted in the different cell cycle phases (EG1, LG1, G1/S, S/G2/M). Samples generated from two independent experiments, 100,000 were used per sample, as described above. Library preparation and sequencing were performed at the Wellcome Trust Sanger Institute next-generation sequencing facility. 8 ATAC-seq libraries were prepared with one of i5 and i7 Nextera tags combination and pooled equimolarly. Sequencing was performed on Illumina HiSeq 2000, 2 X 75bp paired-end reads obtaining more than 60M mapped reads per library. Reads were mapped to the human genome GRCh38 reference assembly and stored as cram files. Duplicates were removed with Samtools rmdup^37^. Cram were converted in bam keeping only reads with mapping score above 5 (q>5) and from chromosome 1-22 and X (H9 is a female line) to remove mitochondrial contamination using Samtools view^37^. From two independent replicates, consistency between replicates was measured using the Irreproducibility Discovery Rate (IDR) pipeline, as described above. Conservative peak set was used for further analyses. Diagnostic plots of fragment size distribution were generated using the R package ATACseqQC^45^. For visualisation purposes, biological replicates were combined as described above to generate normalised bigWig.

Differential accessibility was assessed using the G-test implemented in diffReps^8^ by Dr Pedro Madrigal. This tool takes into account the biological variations within a group of samples and uses that information to enhance the statistical power. For this, we used a sliding window of 600bp, p < 1E-6. Regions with differential binding that did not overlap with previously detected peaks through the IDR pipeline were removed. Genomic regions were ranked by their fold change (FC), and reported as differential only if both the |FC| >1.5 and corrected p-value padj < 1E-6. Footprinting analyses were done by Dr Pedro Madrigal combining two methods: Footprint Log-likelihood Ratio (FLR)^9^ and Wellington^10^, adjusted for ATAC-seq. To get high quality reproducible footprints, motif matches with FLR ≥ 10.0 in each biological replicate were classified as “bound”. Transcription factor footprints that were present in both replicates, for a motif with at least 50 footprints, were considered as final predicted transcription factor binding sites. FLR mean of the replicates was calculated, only based on unique genomic regions (for those palindromic sequences with a footprint reported in each strand, we obtained the mean FLR). The quality of footprint was assessed by calculating the Protection Score, removing scores below 0^46^. Wellington instead uses a “non-motif centric” approach, which considers the imbalance in this strand-specific information to efficiently identify DNA footprints. For this, a relaxed cut-off has been used (-p32 -A -fdr 0.1 --FDR_limit -4 --pv_cutoffs −4 -fp 4,30,1 - fdriter 500—one_dimension) on merged replicates (to increase sequencing depth), in order to achieve a significant overlap with FLR, and a final list generated. A conservative list of footprints was generated by overlap between FLR and Wellington, removing not expressed genes according to RNA-seq data (Fragments Per Kilobase of transcript per Million mapped reads, FPKM > 1.0).

For visualisation of footprints, bigWigs of ATAC-seq data containing Tn5 insertion frequencies have been generated using NucleoATAC^47^ from the bam files (--bam $bamFile --out $outputBasename --cores 16).

### RNA-seq

TRA-1-60+ hESCs were sorted in the different cell cycle phases (EG1, LG1, S/G2/M). From three independent experiments, RNA was extracted using GenEluteTM Mammalian Total RNA Miniprep Kit and On-Column DNase I Digestion Set (both Sigma-Aldrich) and a total of 9 samples fulfilling the submission requirements (0.5-1.0μg total RNA in nuclease free dH2O with A260/A280 and A60/A230 ratios above 1.8) were sent for library preparation and nextgeneration sequencing at CGS (Cambridge Genomic Services). PolyA purified library preparation kit was used, and samples were multiplexed in one HIGH Output NextSeq run Cycle, 2 X 75bp, paired-end reads. A total of 462,569,268 reads evenly distributed across the lanes were obtained (50,547,439 on average per sample). Reads were trimmed using Sickle^48^, an application for trimming FASTQ files based on quality values (Sanger), with a q=20 (quality threshold) and l=30 (length threshold). For the alignment, the transcriptome was built with TopHat2^49^ (v2.1.0), based on Bowtie2^50^ (v2.2.6) using as reference genome the human GRCh38.p6 and GTF annotation from Ensembl (http://ftp.ensembl.org/pub/release-83/gtf/homo_sapiens/). Alignment was performed using TopHat2 with standard parameters. Using Samtools view^37^, reads with mapping quality > 10 were kept for further analyses. Data were quantitated and differential expression analysis was performed using SeqMonk^51^. In sum, data were quantitated using the RNA-seq pipeline, and differential expression analysis was performed using the Intensity Difference Test and the R package DESeq2^13^ (for this, uncorrected raw read counts were used), generating a list of transcripts passing the filter, with a p-value cutoff < 0.05. Hierarchial clustering plots (r > 0.7). Gene Ontology analyses were performed using the Enrichment analysis tool by the Gene Ontology Consortium (Panther)^52^, and validated using Enrich^53^.

### Data availability

The datasets generated during this study are available at ArrayExpress under the following accession numbers: E-MTAB-8347 (RNA-seq), E-MTAB-8348 (ATAC-seq), E-MTAB-8358 (ChIP-seq). Login details for reviewers available.

## Supplementary Information Figure legends

**Supplementary Information Figure 1.**
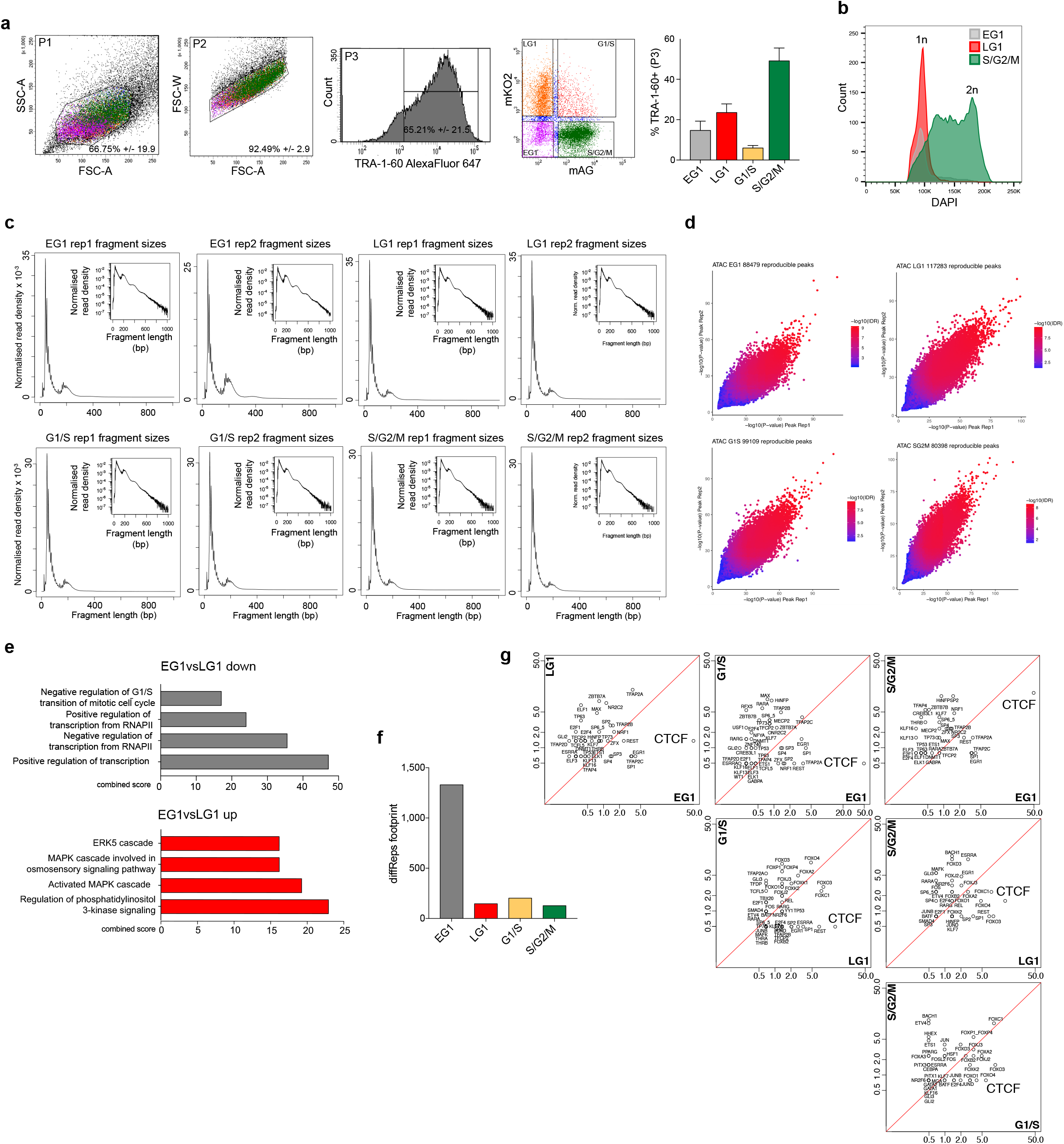
Chromatin accessibility and CTCF footprints during cell cycle progression. **(a)** Gating strategy for FUCCI hESCs: P1 cells are size selected for forward scatter area (FSC-A) and side scatter area (SSC-A); P2, doublets are removed (FSC-A, FSC-W, width); P3 TRA-1-60+ cells are selected and sorted in the four different cell cycle phases according to the FUCCI reporter: EG1 for double negative (mKO2-/mAG-), LG1 for red-mKO2+, G1/S transition for double positive (yellow-mKO2+/mAG+), and S/G2/M for green-mAG+. For all events, average and standard deviation are reported with n=10. **(b)** DNA content analysis by DAPI staining of the different cell cycle phases in FUCCI hESCs. To validate the reporters, EG1 and LG1 show 1n (in grey and red), S/G2/M 2n (in green). **(c)** ATAC-seq fragment size density plot for the 4 sequenced libraries, EG1, LG1, G1/S, and S/G2/M sorted cells. Inset histograms (log-transformed) show periodicity to four nucleosomes. **(c)** IDR (Irreproducible Discovery Rate) plots reporting reproducible ATAC-seq peaks between two replicates in EG1, LG1, G1/S, S/G2/M. IDR threshold < 5%. **(e)** Gene ontology for Biological Process analysis of differentially accessible peaks in EG1vsLG1 down, EG1vsLG1 up, (diffReps, Fig. 1f). Combined score is calculated by multiplying the unadjusted p-values with the z-score. **(f)** Number of predicted footprints overlapping with dynamic regions (ATAC-diffReps) in the different cell cycle phases. Final footprints were obtained combining FLR (Yardimici et al., 2014) and Wellington (Piper et al., 2013) pipelines (combining list with Footprint Likelihood Ratio (FLR) ≥10.0, Protection Score > 0). **(g)** Pairwise comparisons of all cell cycle phases, reporting percentages of footprints for specific transcription factors, highlighting CTCF abundance in all comparison. Log scale axis.

**Supplementary Information Figure 2.**
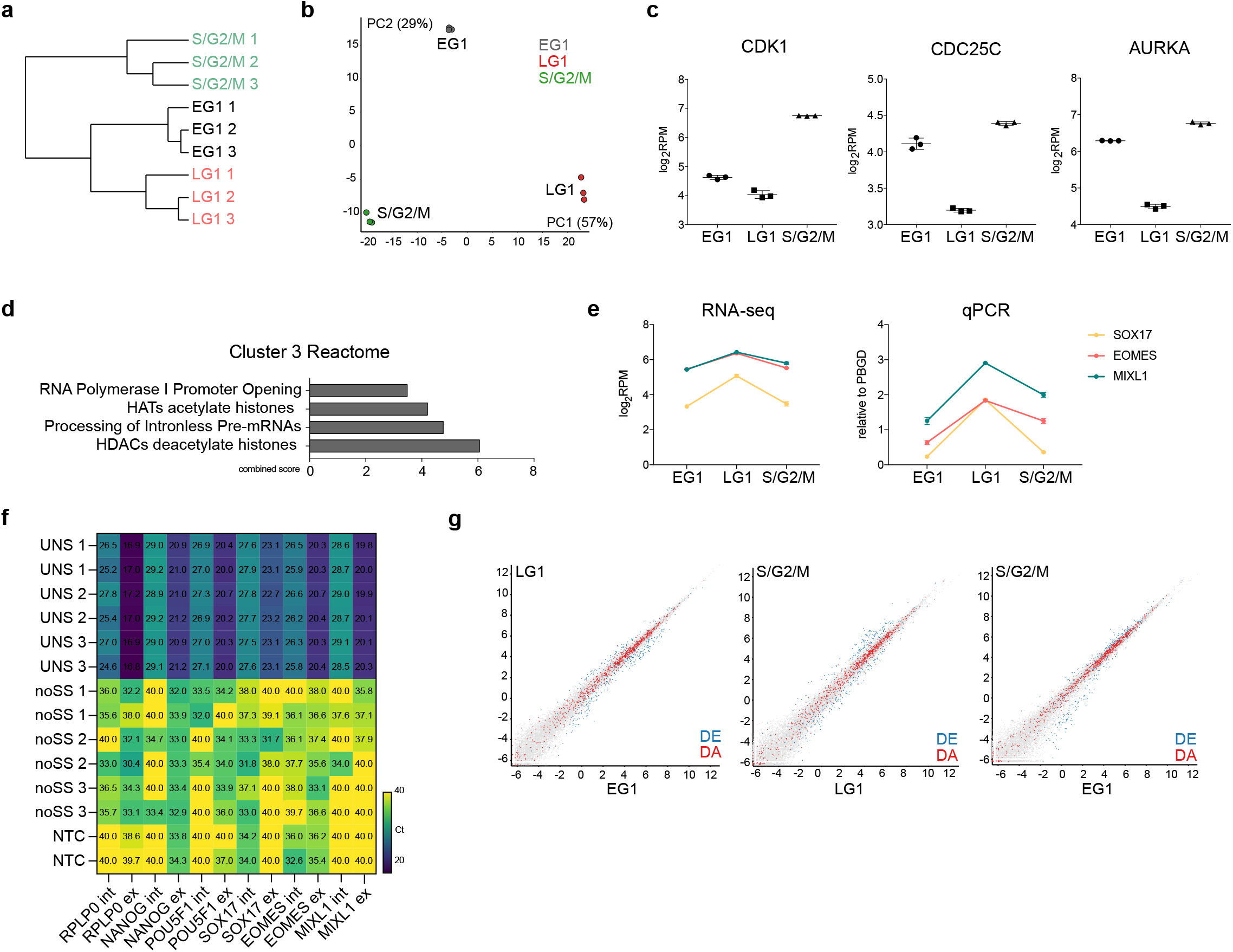
Transcriptome changes during cell cycle progression. **(a)** Datastore Tree clustering of different samples/replicates based on Pearson correlation. **(b)** PCA clustering of FUCCI hESCs sorted in EG1, LG1, S/G2/M. Each dot represents a biological replicate, Principal component contribution reported in brackets. **(c)** Examples of differentially expressed cell cycle regulators *CDK1, CDC25C, AURKA*, up-regulated in the S/G2/M phase (p-value cut-off < 0.05). **(d)** Gene ontology for Reactome Pathway analysis for Cluster 3 (Fig. 2c). Combined score is calculated by multiplying the unadjusted p-values with the z-score. **(e)** RNA-seq log_2_RPM values for the developmental regulators *SOX17, EOMES*, and *MIXL1*, validated by RT-qPCR (values relative to housekeeping gene *PBGD*). **(f)** Heatmap reporting RT-qPCR data of unsorted FUCCI hESCs including non-retrotranscribed RNAs (noSS) as a negative control for genomic contamination in exonic vs intronic data (Fig. 2g). Data expressed as ct values of 3 biological replicates, in technical duplicate. **(g)** Scatter plots reporting pairwise comparison of normalised gene expression by RNA-seq in the different cell cycle phases. Highlighted are differentially expressed genes (DESeq2, DE, in blue) and differentially accessible regions (Fig. 1f, DA, in red), each dot represents one gene.

**Supplementary Information Figure 3.**
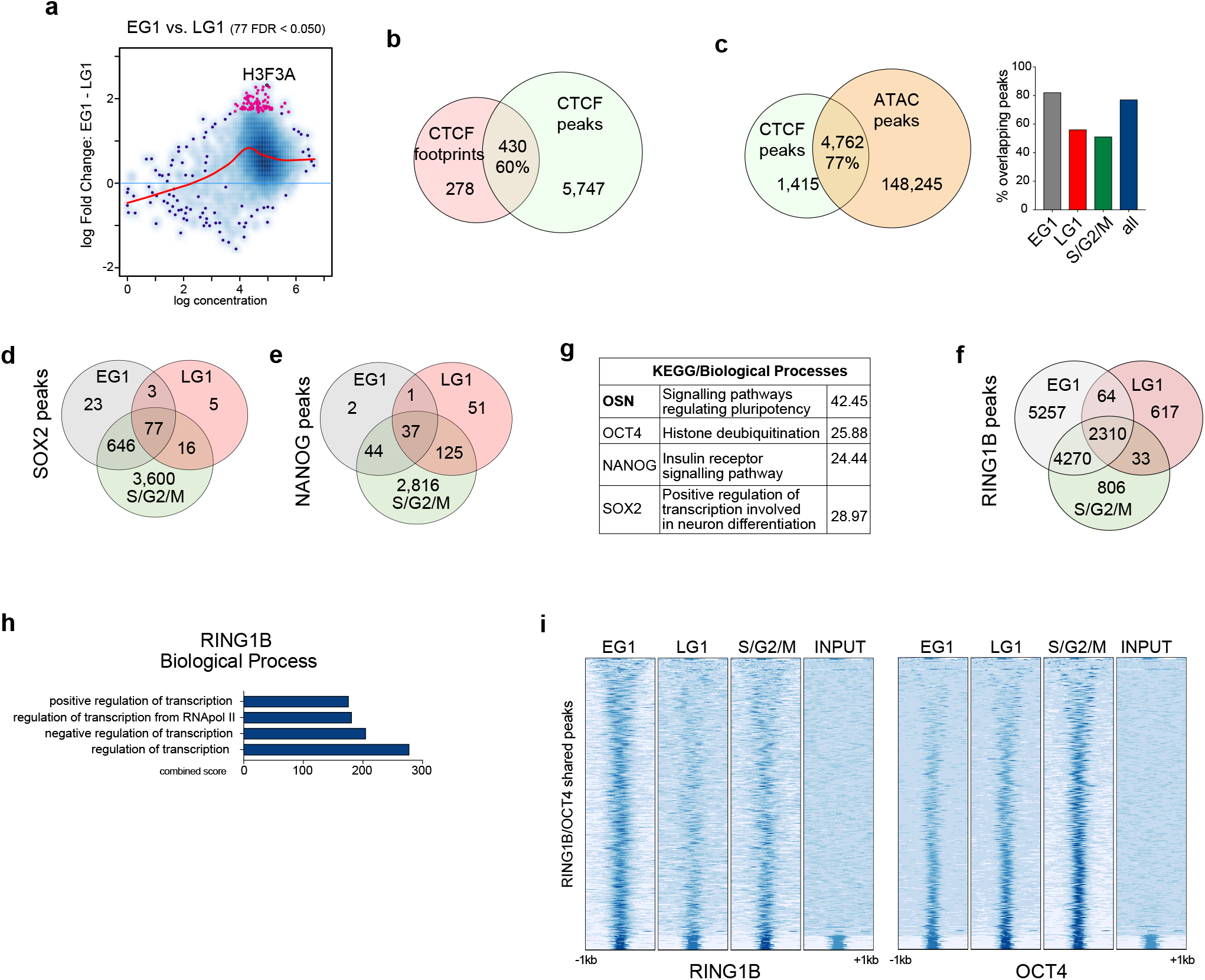
Transcription factors binding dynamics during cell cycle. **(a)** MA plot of CTCF binding sites between EG1 and LG1 phases of the cell cycle in sorted FUCCI hESCs. Each point represents a binding site, in magenta are highlighted the differentially bound sites. **(b)** Venn diagram reporting the overlap of CTCF footprints predicted by ATAC-seq and CTCF ChIP-seq peaks. 60% represent the percentage of footprints overlapping with CTCF ChIP-seq peaks. **(c)** Venn diagram and cell cycle specific quantification of the overlap of CTCF ChIP-seq peaks and ATAC-seq peaks. 77% represent the overall percentage of CTCF ChIP-seq peaks with ATAC-seq peaks. **(d)** Venn diagrams showing peaks overlap for SOX2 and **(e)** NANOG ChIP-seq in EG1, LG1, S/G2/M. **(e)** Gene ontology for KEGG Pathways and Biological Process of OCT4/SOX2/NANOG shared binding sites (OSN) and sites bound by single factors. Values represent the combined score, which is calculated by multiplying the unadjusted p-values with the z-score. **(f)** Venn diagrams showing peaks overlap for RING1B ChIP-seq in EG1, LG1, S/G2/M. **(h)** Gene ontology for Biological Process of RING1B binding sites. Combined score is calculated by multiplying the unadjusted p-values with the z-score. **(i)** Heatmaps displaying normalised ChIP-seq signal ±1kb around all RING1B/OCT4 shared peaks in the different cell cycle phases for both RING1B and OCT4 ChIP-seq, compared to relative INPUTs.

**Supplementary Information Figure 4.**
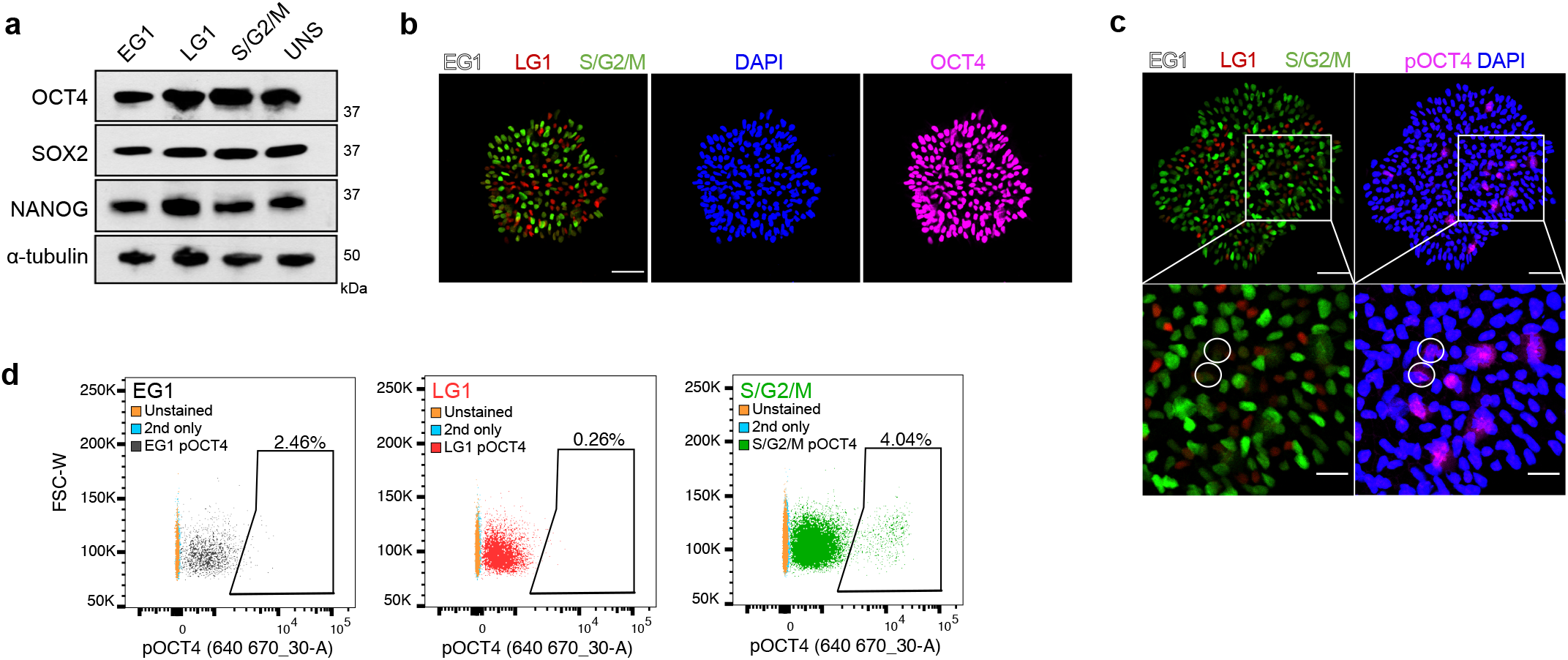
Post-translational phosphorylation of OCT4. **(a)** Additional western-blot analysis for OCT4, SOX and NANOG in EG1, LG1, and S/G2/M, plus unsorted (UNS) control (α-tubulin as loading control). **(b)** Control immunofluorescence for OCT4 (magenta) and nuclear staining (DAPI, blue) in FUCCI hESCs (S/G2/M in green-mAG, LG1 in red-mKO2, EG1 no fluorescence). Scale bar 50μm. **(c)** Additional immunofluorescence for phospho-OCT4 Ser236 (pOCT4, magenta) and nuclear staining (DAPI, blue) in FUCCI hESCs (S/G2/M in green-mAG, LG1 in red-mKO2, EG1 no fluorescence). Bottom panel reports zoom-ed in area from top panel, cells circled are in the EG1 phase. Scale bars: top panel 50μm, bottom panels 20μm. **(d)** Flow cytometry plots of FUCCI hESCs positive for pOCT4 divided by cell cycle phase (EG1 grey, LG1 red, S/G2/M green). Unstained cells and secondary (2nd only) reported for all as negative controls. Quantification in Fig. 4i.

**Supplementary Information Figure 5.**
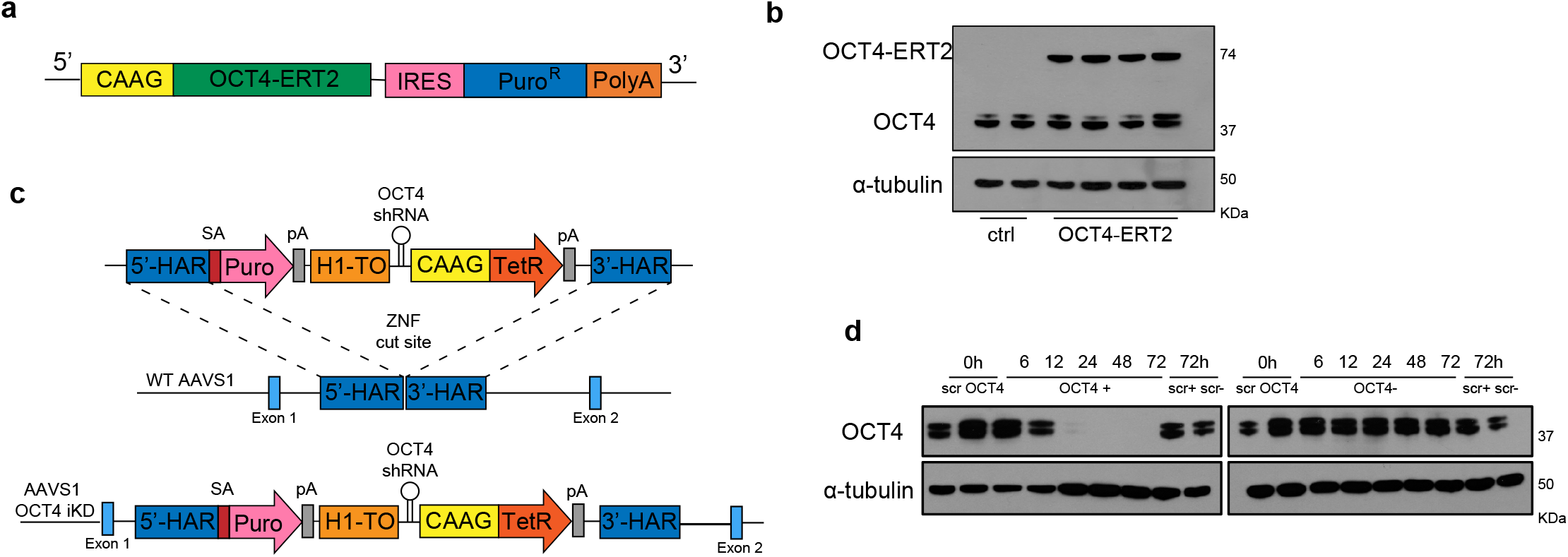
Cell cycle specific OCT4 overexpression and knock-down. **(a)** The modified fragment of the Estrogen receptor (ERT2) is fused to the *OCT4* gene, which is retained in the cytoplasm. In presence of an Estrogen receptor antagonist, such as 4-Hydroxytamoxifen (4OHT), the fusion protein binds to it and relocates into the nucleus. CAAG: CMV early enhancer, chicken β-actin and rabbit β-globin hybrid promoter; IRES: internal ribosome entry site; PuroR: Puromycin resistance. **(b)** OCT4 WB for 2 control lines (lane 1-2) and 4 OCT4 ERT2 lines (lanes 3-6), wild-type OCT4 band at 39 kDa and the OCT4 ERT2 fusion-protein band at 74 kDa. α-tubulin as loading control. **(c)** Single step optimised inducible knock-down^27^. AAVS1 locus targeting, with OCT4 shRNA under the control of a tetracycline inducible promoter. ZFN: zinc-finger nucleases; 5’-HAR/3’-HAR: upstream/downstream homology arm; H1-TO: Tetracycline-inducible H1 Pol III promoter carrying one tet operon after the TATA box; CAAG: CMV early enhancer, chicken β-actin and rabbit β-globin hybrid promoter; TetR: Tetracycline-sensitive repressor protein. **(d)** OCT4 knock-down WB at 0h, 6h, 12h, 24h, 48h, 72h after TET induction (+TET), compared to -TET control. α-tubulin as loading control.

